# Environmental regulation of gene expression mediated by Long non-coding RNAs

**DOI:** 10.1101/2022.06.17.496488

**Authors:** Jingliang Kang, Arthur Chung, Sneha Suresh, Lucrezia L. Bonzi, Jade M. Sourisse, Sandra Ramirez, Daniele Romeo, Natalia Petit-Marty, Cinta Pegueroles, Celia Schunter

## Abstract

The majority of the transcribed genome does not have coding potential but is composed of non-coding transcripts that are involved in transcriptional and post-transcriptional regulation of protein-coding genes. Regulation of gene expression is important in determining the response of organisms to changes in the environment, and therefore their persistence as population or species under global change. However, long non-coding RNAs (lncRNAs) are scarcely studied especially in non-model organisms due to the lack of a reliable pipeline for their accurate identification and annotation. Here, we present a pipeline which uses a combination of alignment-dependent and independent methods for the identification of conserved and species-specific lncRNAs from RNA-Seq data. Validation of this pipeline was performed using existing RNA-Seq data from *Acanthochromis polyacanthus* brain tissue, identifying a total of 4,728 lncRNAs across the genome, the majority of which (3,272) are intergenic. To investigate the possible implications of these intergenic lncRNAs (lincRNAs), we estimated the expression changes of lincRNAs and coding genes in response to ocean acidification. We found lincRNAs which neighbour or possibly trans-regulate differentially expressed coding genes related to pH regulation, neural signal transduction and ion transport, which are known to be important in the response to ocean acidification in fish. Overall, this pipeline enables the use of existing RNA sequencing data to reveal additional underlying molecular mechanisms involved in the response to environmental changes by integrating the study of lncRNAs with gene expression.

## INTRODUCTION

Among pervasive genomic regions that can be transcribed, some encode long non-coding RNAs (lncRNAs), defined as RNAs longer than 200 nucleotides that are not translated into functional proteins (Statello et al., 2021). LncRNAs represent a highly heterogeneous class of transcripts mainly transcribed by RNA polymerase II and, usually, inefficiently spliced. While some lncRNAs remain in the nucleus, others are polyadenylated and exported to the cytoplasm (Statello et al., 2021). Although initially considered to be products of transcriptional noise or spurious transcription (Struhl, 2007), lncRNAs are now known to play critical roles in diverse biological processes, including DNA repair, proliferation, and embryonic development (Fernandes et al., 2019; Li et al., 2019; Vance & Ponting, 2014). LncRNAs are involved at many levels, including chromatin modifications, pre-transcription, transcription, and post-transcription through regulation of the associated gene expression (Gardini & Shiekhattar, 2015; Kornfeld & Brüning, 2014; Necsulea et al., 2014; Ulitsky, 2016; H. Xu et al., 2019; Zhu et al., 2013). In addition, lncRNAs are involved in specific physiological processes and gene dysfunction and can be used as biomarkers of certain diseases (Beck et al., 2018; Fernandes et al., 2019; Jiang et al., 2016).

LncRNAs are not as conserved as protein-coding sequences due to their fast turn-over (Lopez-Ezquerra et al., 2017; Pegueroles et al., 2019), and they tend to be cell and tissue specific (Cabili et al., 2015; Derrien et al., 2012). In addition, many lncRNAs are lowly expressed (Guo et al., 2020; Hezroni et al., 2015; Necsulea et al., 2014; Quinn et al., 2016). As such, despite the growing interest in lncRNAs and the availability of RNA sequencing data in many species, lncRNAs remain unexplored in most species, particularly in non-model species. To date, several bioinformatic tools have been developed to detect lncRNAs by computing coding potential score (CPS) of transcripts, either by using sequence alignments (alignment-dependent) or detecting intrinsic features of the input RNA sequences (alignment-free). In the alignment-dependent methods, such as PhyloCSF (Lin et al., 2011) and CPC (Kong et al., 2007), sequences are either aligned between species or to protein databases, which may be biased toward misclassifying species-specific or lowly conserved coding and non-coding transcripts. In contrast, although the alignment-free methods like CPAT (Wang et al., 2013) and FEELnc (Wucher et al., 2017) are useful to discriminate species-specific lncRNA, these tools use different intrinsic features (such as the length and integrity of the longest open reading frame) of the input RNA sequences, which can result in differences in detecting lncRNAs.

Nevertheless, most studies related to lncRNA identification used only one method (Boltaña et al., 2016; Dettleff et al., 2020; Mu et al., 2016; Paneru et al., 2018; D. Quan et al., 2020; Ren et al., 2020; J. Xu et al., 2019). Hence, a pipeline combining both alignment-dependent and alignment-free strategies is essential to detect conserved and species-specific lncRNAs.

Environmental perturbations produce molecular responses in organisms needed to maintain cellular homeostasis and function, and lncRNAs may play a role in some of these responses by regulating gene expression (Paneru et al., 2018; Sarangdhar et al., 2018) through cis- and trans-regulation (Gil & Ulitsky, 2020; Luo et al., 2019; Nadal-Ribelles et al., 2014; D. Quan et al., 2020). A large proportion of lncRNAs with functional implications are in fact cis-regulatory and hence affect the regulation of the neighbouring protein-coding genes allowing for plasticity in gene networks across time (Engreitz et al., 2016; Gil & Ulitsky, 2020). In the model-plant genus *Arabidopsis*, for instance, a lncRNA tightly controls the expression of highly conserved transcription factors that promote cold tolerance in many plant species (Kindgren et al., 2018). In vertebrates, such as fish, lncRNAs identified in rainbow trout *(Oncorhynchus mykiss*) can potentially mediate regulation of the heat stress response (J. Quan et al., 2020) and one lncRNA affects the mucosal immunity of turbot (*Scophthalmus maximus*) in response to bacterial infection (N. Yang et al., 2020). The involvement of lncRNAs in the regulation of the response to environmental change reveals its importance as a potential regulator of (phenotypic) plasticity and in turn adaptive potential in rapidly changing environments. Hence, there is a need for more studies to clarify the role of lncRNAs in the molecular response of species to changing environment.

To this end, we propose a comprehensive roadmap and guideline for the *de novo* identification of lncRNAs and use a case study to evaluate the involvement of lncRNAs and related coding genes in the response to an environmental change. For this, we use the spiny chromis *Acanthochromis polyacanthus*, a common coral reef fish in the Western Pacific, extensively studied for fish acclimation processes to environmental changes in the last decade. Generous RNA sequencing has been performed on this species (Bernal et al., 2022; Kang et al., 2022; Schunter et al., 2021, 2016) to interpret its transcriptional responses to ocean environmental changes. Despite the abundance of transcriptional information nothing is known about the potential of lncRNAs acting as modulators of gene expression in response to environmental stress. This extensive knowledge on the transcriptional processes to ocean acidification in *A. polyacanthus* allows for a broad evaluation of the involvement of lncRNAs and provides a guideline for future studies also in other systems. As lncRNAs can tune neighbouring coding gene expression by directly affecting nuclear architecture or indirectly affecting their transcription or translation (Ransohoff et al., 2018) we aim to combine the study of lncRNAs with coding gene expression to evaluate the transcriptional and epigenetic factors that define the biological response of organisms to environmental changes, providing information on the adaptive potential of the species.

## METHODS

### Discovery and Annotation of lncRNAs

To identify lncRNAs in the *Acanthochromis polyacanthus* genome, 226 RNA-seq samples were compiled from published studies investigating the effect of elevated CO_2_ on the brain transcriptome of *A. polyacanthus* (Kang et al., 2022; Schunter et al., 2018, 2016; PRJNA658203). This allows for a comprehensive annotation across the genome of lncRNAs associated with the exposure to elevated CO_2_ in laboratory and natural settings. We processed all samples to identify a reliable set of lncRNAs across the genome (Figure 1 & S1). The RNAseq reads are paired-end and non-strand specific, as most RNA sequencing datasets in molecular ecological studies. In our pipeline, the raw reads quality was first assessed with FASTQC v0.11.9 (Andrews, 2010), and then adapter sequences and low-quality sections of reads were removed by Trimmomatic v0.39 (Bolger et al., 2014) using the following parameters: “ILLUMINACLIP:2:30:10 LEADING:4 TRAILING:3 SLIDINGWINDOW:4:20 MINLEN:40”. The resulting clean reads were aligned to the reference genome of *A. polyacanthus* (NCBI database, accession number: ASM210954v1) with HISAT2 v2.1.0 (Kim et al., 2019) using default parameters and with “*--known-splicesite-infile”* to provide known splice sites.

**Figure 1:**
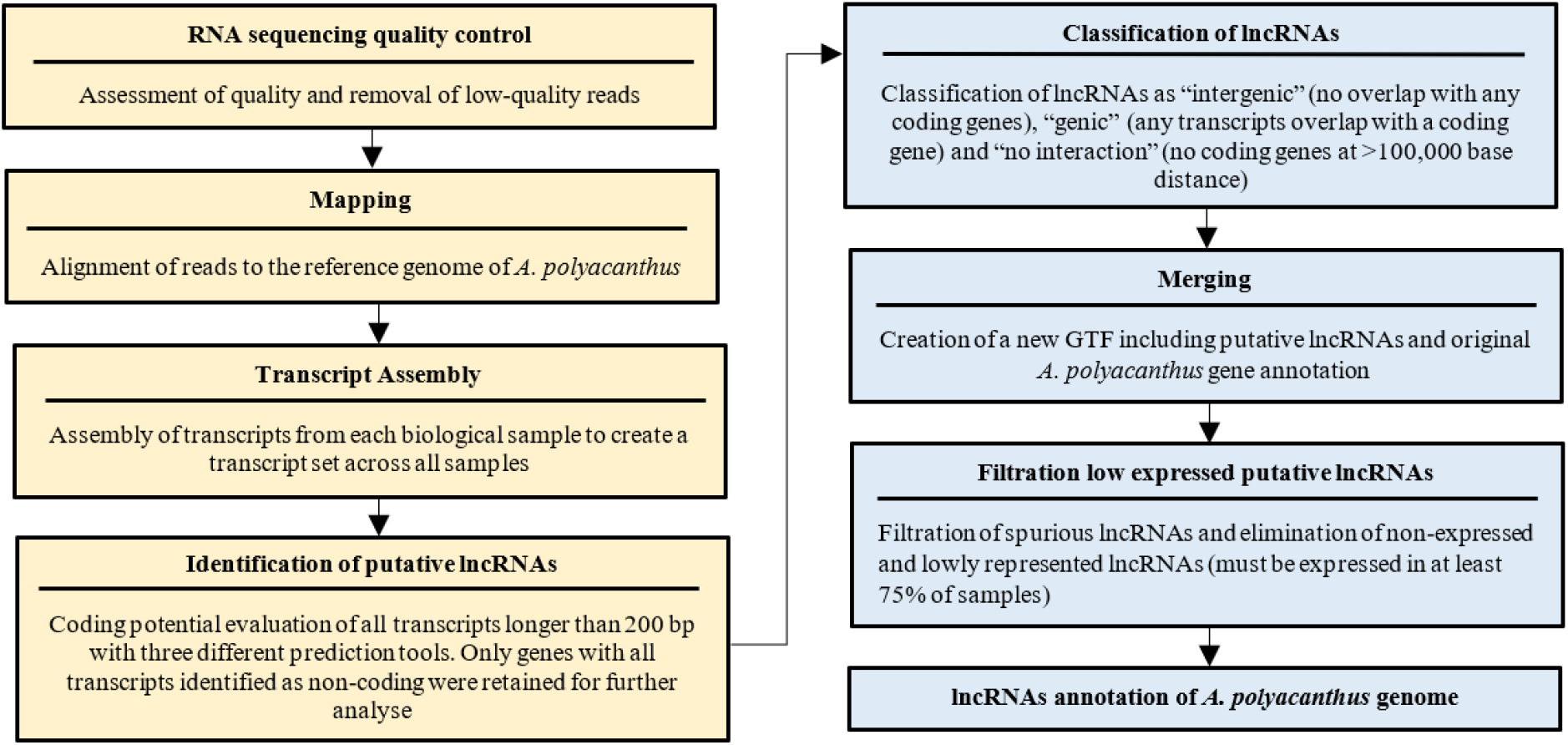
Flowchart illustrating the process for long non-coding RNA (lncRNA) identification in Acanthochromis polyacanthus genome. In yellow are the steps for the initial identification of putative lncRNAs, while in blue are the post-processing analyses for the creation of the final high confidence lncRNA set.

StringTie v2.1.5 (M. Pertea et al., 2015) was then applied to assemble transcripts. All resulting transcripts among the 226 fish samples were merged for a unified set of non-redundant transcripts (M. Pertea et al., 2016).

To estimate the coding potential of all transcripts, we extracted the transcript sequences from the *A. polyacanthus* genome sequence using GffRead v0.12.7 (G. Pertea & Pertea, 2020) with the parameters *“-w, -g”*. Three tools, including one alignment-dependent (CPC v0.9-r2, Kong et al., 2007) and two alignment-free methods (CPAT v1.2.4, Wang et al., 2013; FEELnc v0.2, Wucher et al., 2017), were used to calculate the coding potential of all transcripts. CPC v0.9-r2 (Kong et al., 2007) was used with default parameters to estimate coding potential based on sequence similarity between transcripts and Uniref90 protein database, and the transcripts with a coding potential score < 0 were considered as putative lncRNAs. For the two alignment-free tools, CPAT v1.2.4 (Wang et al., 2013) was run with default parameters using the coding mRNA sequences of zebrafish as training model, and transcripts with a coding probability < 0.38 were considered as putative lncRNAs, which is the cut-off value suggested by CPAT (Wang et al., 2013). The second alignment-free tool we used was FEELnc v0.2 (Wucher et al., 2017), which annotates lncRNAs based on a Random Forest model trained with general features such as multi k-mer frequencies and relaxed open reading frames. We ran the program with FEELnc_filter.pl to filter out spurious coding transcripts using “-b transcript_biotype=protein_coding, --monoex=-1”, and then FEELnc_codpot.pl with default parameters to compute the coding potential score for each transcript by setting the mode of the lncRNA simulation to intergenic. The putative lncRNAs were then identified based on the best coding potential cutoff of receiver operating characteristic curve plot (Sing et al., 2005). For all three methods, only genes with all transcripts identified as non-coding were retained for further analyses.

For a more conservative approach in avoiding spurious lncRNA annotations we only retained the transcripts that were identified as lncRNAs by at least two tools. Using a window size 100 kilobase (kb) (Wucher et al., 2017), these putative lncRNAs were categorized by computing interactions with their proximal coding transcripts using default parameters of FEELnc_classifier.pl (Wucher et al., 2017). A lncRNA is classified as “intergenic” (i.e., lincRNA) if all transcripts of this lncRNA have no location overlap with any neighbouring. Due to the fact that intergenic lncRNA (lincRNA) gene expression patterns, sequence conservation and perturbation outcomes are easier to interpret than those of transcripts from other lncRNAs, for instance overlapping with coding genes (“genic” lncRNAs; Ulitsky & Bartel, 2013), only lncRNA genes classified as intergenic were used in further analyses to investigate their potential involvement in the response to ocean acidification in fish.

The expression levels of the putative lincRNAs and mRNAs were quantified using FeatureCounts v2.0.0 (Liao et al., 2014) allowing for multi-mapped reads to be counted fractionally. Only transcripts with reads number > 0 in at least 75% of the samples were retained for further analysis. The read numbers of remaining transcripts were normalized using DESeq2 v1.32.0 (Love et al., 2014) and lincRNAs with a normalised expression < 1 in 10% samples were removed to obtain the final list of candidate lincRNA genes. This provided us with a final set of high confidence lincRNAs in both location and expression. LincRNAs with more than 500 normalized reads were considered as highly expressed lincRNAs.

### Case study: Expression patterns of lincRNAs in fish living in CO_2_ seeps

Using RNA sequencing data from wild *A. polyacanthus* samples collected in Papua New Guinea (Kang et al., 2022; NCBI Bioproject PRJNA691990), we performed a case study to evaluate the expression changes of lincRNAs in different environmental conditions. Seven brain samples were collected from fish from a coral reef situated in a naturally bubbling CO_2_ seep with elevated CO_2_ levels close to the predicted levels for the end of this century due to ocean acidification (pH= 7.77, *p*CO_2_= 843 μatm; IPCC, 2022). Further eleven fish were sampled from an adjacent control reef approximately 500 m away from the CO_2_ seep (pH= 8.01, *p*CO_2_= 443 μatm; Kang et al., 2022). There was no significant difference in temperature and salinity between the CO_2_ seep and the control site (Fabricius et al., 2011) allowing for the evaluation of effects of long-term ocean acidification conditions on fish.

To evaluate differential gene expression of lincRNAs and coding genes between fish living at different CO_2_ levels, we performed differential gene expression analysis using DESeq2 v1.34 (Love et al., 2014) in R v.3.5.1. Between samples from control and CO_2_ seep, lincRNAs and coding genes were considered as differentially expressed (DE) with an FDR adjusted p-value ≤ 0.05, and the average of the normalized count values (basemean) ≥10 as well as Log2FoldChange ≥0.3. A principal component analysis (PCA) was performed using the log 2-fold normalized expression of the samples from the two different CO_2_ level sites.

To identify lincRNAs and coding gene co-expression modules with significant correlation with CO_2_ levels, a weighted gene co-expression network analysis (WGCNA; Langfelder & Horvath, 2008) was also applied on our fish samples. The co-expression similarity of gene modules was obtained by selecting a *signed* network adjacency type with a soft thresholding power of 10 calculated according to the scale-free topology criteria. We established the expression similarity between nodes of genes that are co-expressed by calculating the Topological Overlap Measure (TOM). These co-expression networks were grouped into different coloured modules based on their eigengenes values. Pearson correlation was then calculated to evaluate the correlation between gene modules and samples from control and CO_2_ seep. Genes of significantly correlated gene module (p < 0.01) were used in the following analysis.

### Prediction of potential cis- and trans-acting lincRNAs

To investigate the potential role of cis-acting lincRNAs in the response to elevated CO_2_ we focused on lncRNAs which are neighbouring to coding genes with a maximum of 100 kilobase (kb) distance (Wucher et al., 2017). We performed several analyses to investigate a) neighbouring coding genes of highly expressed lincRNAs, b) neighbouring differentially expressed (DE) coding genes of DE lincRNAs, and c) neighbouring coding genes of lincRNAs found co-expressed within the same WGCNA module.

For potential trans-acting lincRNAs, the reads number of all 226 individuals was normalized by calculating transcripts per kilobase million (TPM), and the genes with TPM ≥ 1 among more than 90% individuals were kept. Based on the TPM values, except for the lincRNAs that have neighbouring coding genes, we estimated the association relationship between each lincRNA-coding gene pair for d) DE lincRNAs and DE coding genes, and e) lincRNAs and coding genes within the same WGCNA module using Spearman’s correlation test (Tsai et al., 2021) in R version 3.6.3. The lincRNAs that showed Spearman’s correlation coefficient |rho| ≥ 0.9 and p value ≤ 0.01 with a coding gene, were considered as potential trans-acting lincRNAs. Such correlated coding genes were used for functional enrichment analysis.

For all subsets, functional enrichment analyses were performed in OmicsBox v2.0.36 (BioBam Bioinformatics, 2019) with all the annotated genes in *A. polyacanthus* genome as reference.

## RESULTS

### LncRNA annotation pipeline

Brain *Acanthochromis polyacanthus* RNA sequencing samples had on average 32,7 million high-quality paired-end reads (Table S1) and on average, 87.5% of these reads mapped to the reference genome (Table S2). The assembly steps resulted in a total of 116,237 transcripts across the genome. We used different algorithms to predict the coding potential score (CPS) of these transcripts. This resulted in 29,127 non-coding transcripts with CPAT, 28,865 with CPC and 11,897 with FEELnc, belonging to 17,434, 7,970 and 3,633 putative long non-coding RNA genes (lncRNAs) respectively. Of these, 9,209 putative lncRNAs were identified by at least two programs (Figure 2A). After filtering out lowly expressed putative lncRNAs, we obtained a final set of 4,728 lncRNAs, consisting of 1,313 genic, 3,272 intergenic and 143 non-neighbouring lncRNAs.

**Figure 2:**
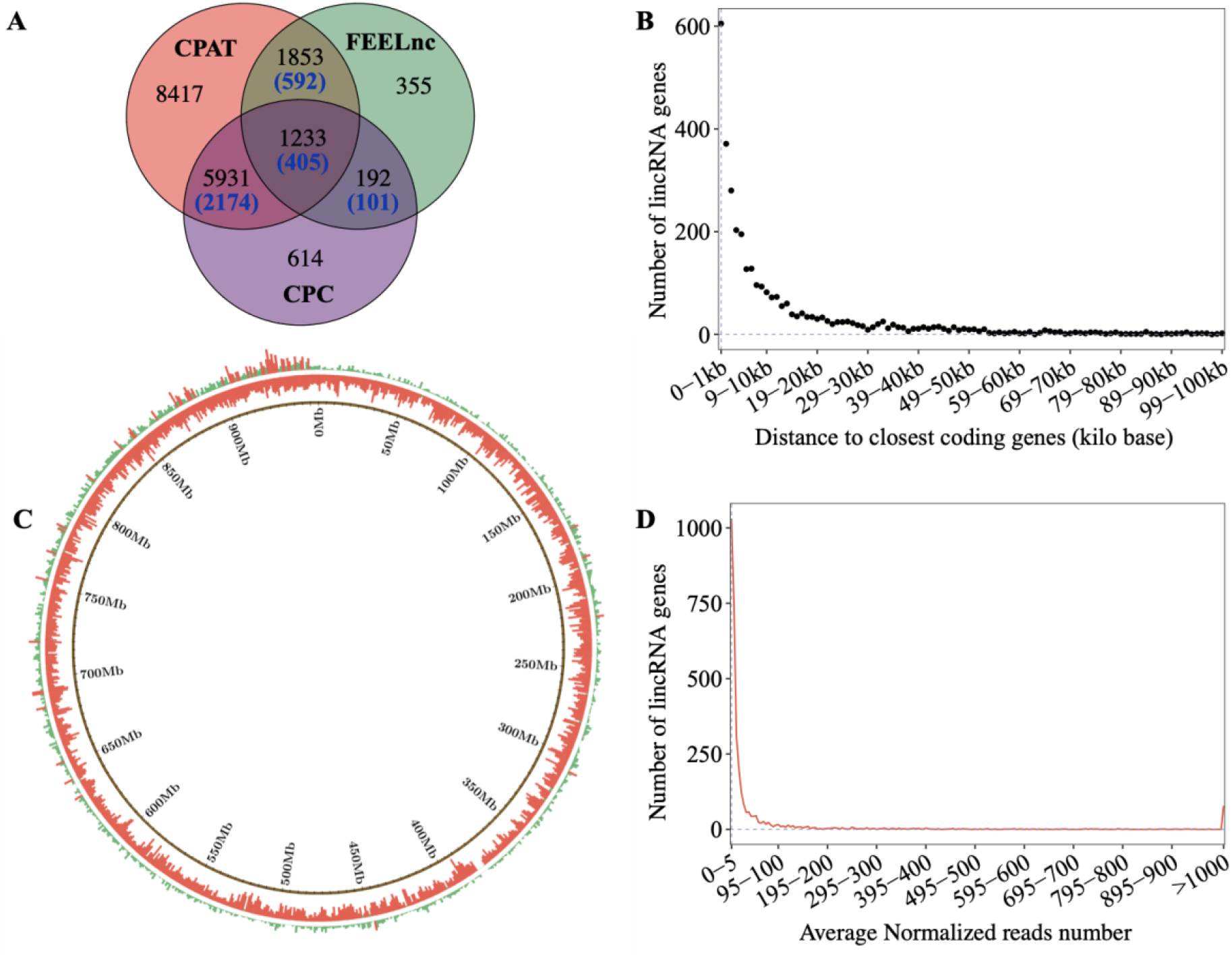
Intergenic lncRNA genes (lincRNAs) along the Acanthochromis polyacanthus genome.**A**. Venn diagram of putative lncRNAs identified by CPAT, CPC and FEELnc programs. Blue numbers in brackets indicate overlapping lincRNAs among the total lncRNAs. **B**. Distance ranges between the identified 3,272 lincRNAs and their neighbouring coding genes. **C**. Distribution of lincRNAs (first outer circle) and coding genes (second outer circle) along the whole genome (concatenating all scaffolds together from longest to shortest). Genome sliding windows are indicated in the inner black circle. Bar height and color are indicative of the number of genes found in each 1Mb sliding window, with red and green bars indicating more and less than 10 lincRNAs or coding genes per window, respectively. **D.** LincRNAs levels of expression across different expression level ranges.

The intergenic lncRNA genes (lincRNAs; Table S3) are significantly shorter in length than coding genes (two-sample wilcoxon rank sum test, p < 2.2e-16) with 1,386 lincRNAs with a length less than 400 nucleotides (Figure S2A). Of all intergenic lncRNA genes (Table S3), 2,143 (65.5%) lincRNAs are monoexonic, while 1,129 (34.5%) lincRNAs have at least two exons (Figure S2B). In comparison, 30,282 coding genes (88.6%) have at least two exons, which are significantly more than lincRNAs (two-sample wilcoxon rank sum test, p < 2.2e-16). The GC content (43.1%) of lincRNAs is significantly lower than coding genes (50.0%, two-sample wilcoxon rank sum test, p < 2.2e-16, Figure S2C). LincRNAs (Table S3) are mostly located within a short distance to their neighbouring coding genes (Figure 2B) with 1,654 (50.6%) of lincRNAs located at a distance smaller than 5kb from their neighbour coding genes. Of these lincRNA, 354 are antisense, 243 sense lincRNAs and 1,057 strand-unknown lincRNAs (Table S3). Among the 354 antisense lincRNAs, four lincRNAs (MSTRG.12220, MSTRG.40264, MSTRG.36665, MSTRG.10872) contain both convergent and divergent transcripts, while 146 and 204 only contained divergent and convergent transcripts, respectively (Figure S3). LincRNAs were dispersed throughout the whole genome (Figure 2C), however, the overall lincRNAs density was significantly lower than in protein coding genes (two-sample wilcoxon rank sum test, *p* < 2.2e-16). Among 955 genomic sliding windows of 1Mb in the *A. polyacanthus* genome, 829 regions included more than one lincRNA and 53 regions included more than ten lincRNAs. Protein coding genes, by contrast, displayed a higher density (934 regions with more than 10 coding genes).

Most lincRNAs exhibited expression with less than 500 normalized reads, and only 125 lincRNAs had elevated expression levels (average normalized reads > 500; Figure 2D; Table S4). The neighbouring coding genes of the 125 most expressed lincRNAs were involved in signal transduction, GABA-A receptor, circadian rhythm, and ion transport (Figure 3A; Table S4). The highest expressed lincRNA (MSTRG.9528; expression > 190,000 normalized reads) neighbours the coding gene peripherin-2 (PRPH2).

**Figure 3:**
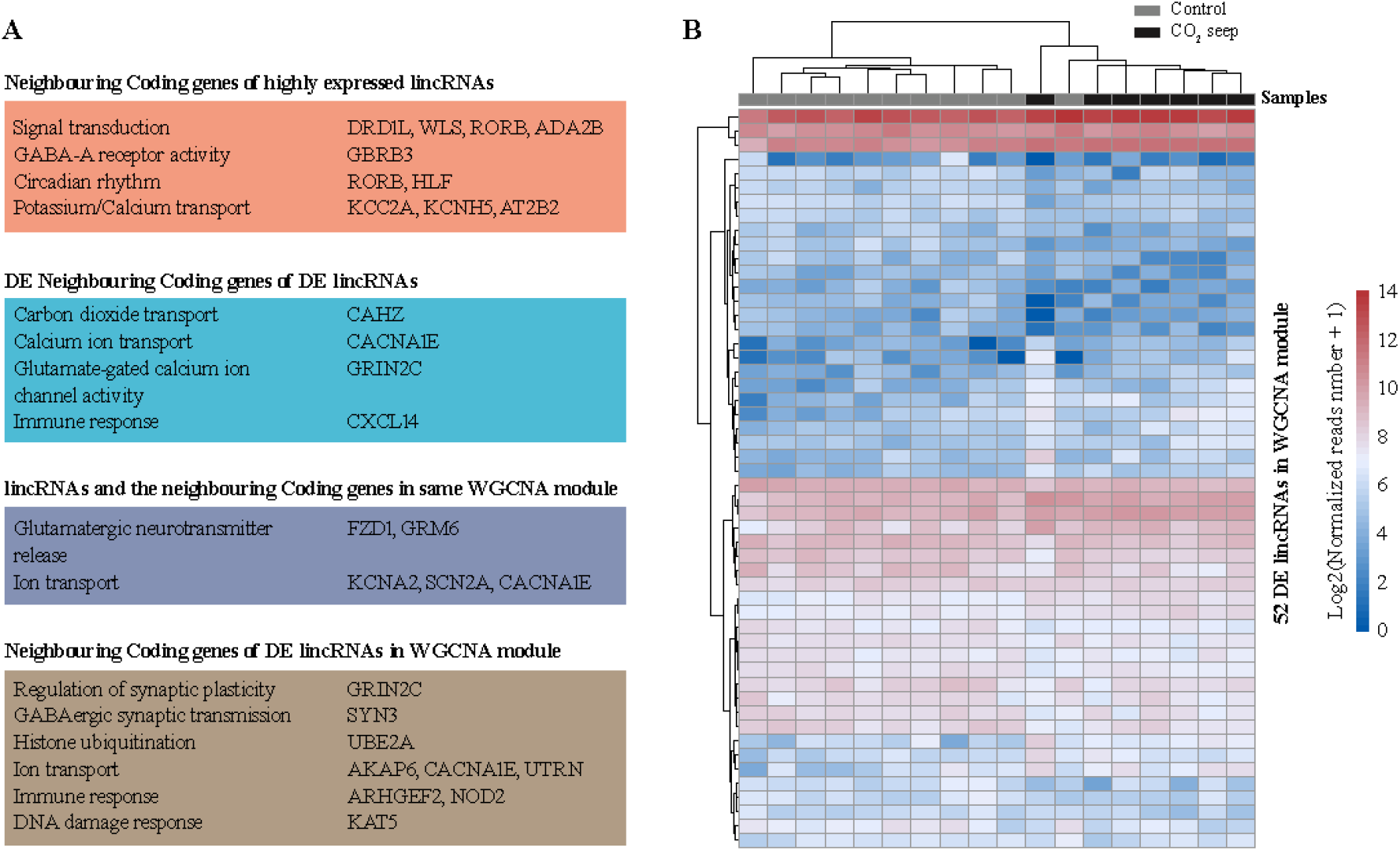
Functions of neighbouring coding genes of lincRNAs (A) and gene expression pattern of 52 differential expressed lincRNAs which were also related to pH level in module turquoise by WGCNA analysis (B).

### Case study: response to ocean acidification in fish

By applying our pipeline to already existing transcriptomic data from wild *A. polyacanthus* collected from a CO_2_ seep and a control reef site in Papua New Guinea, we were able to evaluate the involvement of genomic regulations by lincRNAs in fish brain in response to an environmental factor. Fish from the CO_2_ seep displayed a distinct brain expression pattern, both when the analysis was run using all genes (coding genes and lincRNAs; Figure 4A) as well as only lincRNAs (Figure 4B). Between samples from CO_2_ seep and control site, 3,431 coding genes (Figure S4A) and 97 lincRNAs (Table 1; Table S5) were significantly differentially expressed by DESeq2 v1.34 (Love et al., 2014).

**Figure 4:**
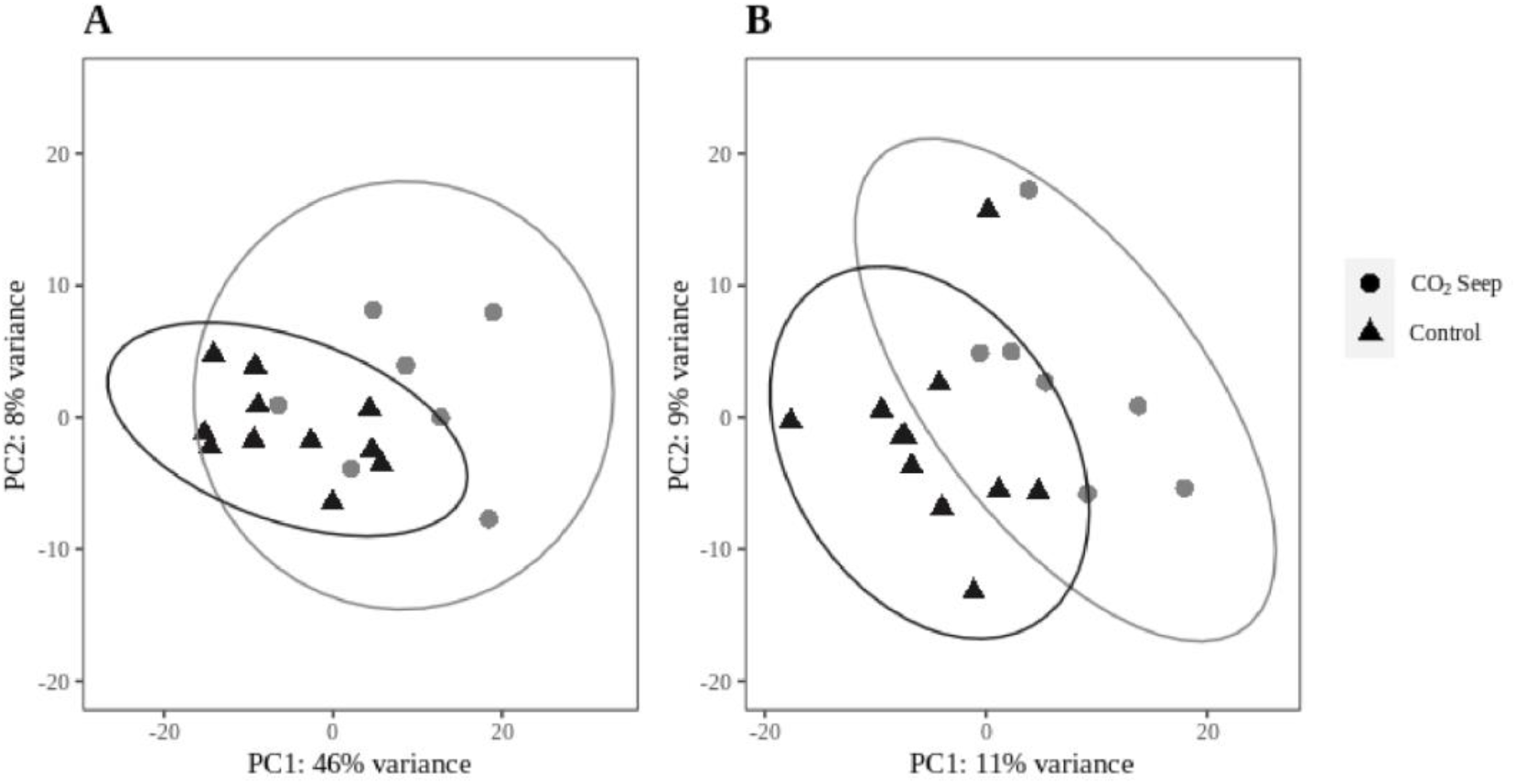
Principal component analyses (PCAs) of log2-fold normalized expression values of (A) lincRNA and coding genes or (B) lincRNAs only in the brain of wild collected Acanthochromis polyacanthus from a CO_2_ seep (circles) and an adjacent control reef (triangles). Ellipse areas represent a 95% confidence level.

**Table 1.** Differentially expressed lincRNAs and coding genes identified through differential expression analysis (DESeq2) and weighted gene co-expression network analysis (WGCNA). In the “lincRNAs with neighbouring coding gene” row are the numbers of lincRNAs for which the neighbouring coding genes were also differentially expressed or found in the same co-expression module.

| Gene type | Differential Expression | Weighted co-expression network |  |  |
| --- | --- | --- | --- | --- |
|  |  | turquoise | sky-blue | dark-green |
| lincRNAs | 97 | 329 | 26 | 6 |
| coding genes | 3431 | 6,259 | 136 | 231 |
| lincRNAs with neighbouring coding gene | 13 | 65 | 0 | 0 |

WGCNA results showed that three modules were significantly correlated with environmental CO_2_ levels (“turquoise”, p = 0.001, Table S7; “sky-blue”, p= 0.009, Table S8; “dark-green”, p= 0.006, Table S9). Turquoise and sky-blue modules were negatively correlated with CO_2_ levels and included 329 lincRNAs and 6,259 coding genes (Table 1; Table S7), and 26 lincRNAs and 136 coding genes (Table 1; Table S8), respectively. Dark green module was conversely positively correlated with CO_2_ levels and consisted of 6 lincRNAs and 231 coding genes (Table 1; Table S9).

### Cis-regulatory roles of lincRNAs in response to elevated CO_2_

Based on the significantly differentially (DE) genes detected by DESeq2, DE coding genes with neighbouring lincRNAs had significantly higher expression changes (Pearson’s Chi-squared test, *p* = 3.425e-05). 3.5% (7/198) of the DE coding genes with neighbouring lincRNA exhibit high expression difference (|log2FoldChange| > 2), whereas only 0.7% (23/3233) of the DE coding genes without neighbouring lincRNA display elevated expression (Figure S4B). Among the neighbouring coding genes of the 97 DE lincRNAs, 13 coding genes also displayed significant differential expression (Table S6). These DE neighbouring coding genes are related to carbon dioxide transport and hypotonic salinity response (CAHZ: carbonic anhydrase), calcium ion transport (CACNA1E: calcium voltage-gated channel subunit alpha1 E), glutamate-gated calcium ion channel activity (GRIN2C: glutamate ionotropic receptor NMDA type subunit 2C) or immune system response (CXCL14: C-X-C motif chemokine 14; Figure 3A).

Among the significantly correlated co-expression networks, only the turquoise module was found with 68 lincRNAs whose neighbouring coding genes were also included in the same co-expression module (Tables S7 & S10). The functional implications of these neighbouring coding genes show involvement in, for instance, glutamatergic neurotransmitter release (FZD1: frizzled class receptor 1; GRM6: glutamate metabotropic receptor 6) and ion transport (KCNA2: potassium voltage-gated channel subfamily A member 2; SCN2A: sodium voltage-gated channel alpha subunit 2; CACNA1E: calcium voltage-gated channel subunit alpha1 E; Figure 3A).

Among the lincRNAs in WGCNA modules, turquoise module showed 52 lincRNAs which were also significantly differentially expressed by DESeq2 (Figure 3; Tables S7 & S11). The neighbouring coding genes of these lincRNAs were related to the regulation of synaptic plasticity (GRIN2C: glutamate receptor ionotropic NMDA 2C), modulation on GABAergic synaptic transmission (SYN3: synapsin-3), histone ubiquitination (UBE2A: ubiquitin-conjugating enzyme E2 A), regulation of ion transport (AKAP6: A-kinase anchor protein 6; CACNA1E: calcium voltage-gated channel subunit alpha1 E; UTRN: utrophin), innate immune response (ARHGEF2: rho guanine nucleotide exchange factor 2; NOD2: nucleotide-binding oligomerization domain-containing protein 2) and DNA damage response (KAT5: histone acetyltransferase KAT5; Figure 3A). Of the other two significantly correlated modules, sky-blue and dark green, only two and one lincRNAs, respectively, were also found to be differentially expressed (Table S8 & S9).

### Regulatory roles of potential trans-acting lincRNAs in response to elevated CO_2_

Between DE lincRNAs and DE coding genes, 148 lincRNA-coding gene pairs exhibited high and significant correlation with |rho| ≥ 0.9 and p value ≤ 0.01, which include 14 lincRNAs and 129 coding genes (Table 2). With the co-expression networks, only the turquoise module revealed 219 lincRNA-coding gene pairs with high and significant correlation made of 14 lincRNAs and 129 coding genes (Table 2). In summary, 21 lincRNAs may trans-regulate 186 coding genes (Table S12), including genes involved in ion transport (KCMA1: Calcium-activated potassium channel subunit alpha-1; SCN8A: Sodium channel type 8 subunit alpha; CACNA1A: Voltage-dependent P Q-type calcium channel subunit alpha-1A; RYR3: Ryanodine receptor 3; NALCN: Sodium leak channel non-selective), glutamate receptor activity (GRM4: Glutamate receptor 4; NMDE1: Glutamate receptor NMDA 2A), and immune response (BCL11A: B-cell lymphoma leukemia 11A; BCL11B: B-cell lymphoma leukemia 11B; BCL9: B-cell CLL lymphoma 9).

**Table 2.** Significantly correlated pairs of lincRNA and coding genes, which were differentially expressed (DE) lincRNAs and DE coding genes from DESeq2, and lincRNAs and coding genes in the same modules from the weighted correlated co-expression network analysis.

|  | LincRNA-coding gene pairs | LincRNAs | Coding genes |
| --- | --- | --- | --- |
| DESeq2 | 148 | 14 | 129 |
| WGCNA | 219 | 14 | 129 |

## DISCUSSION

### De novo *lncRNAs discovery pipeline*

The boost in RNA sequencing studies including a large variety of non-model organisms provides an excellent resource that could be (re-)used to obtain a more complete picture of lncRNAs regulations on coding genes. Here we present a solid pipeline that can make use of already existing data to detect conserved as well as species-specific lncRNAs. This pipeline can be easily applied to any species with an available reference genome. The number of published genomes has been increasing drastically the last few years even in non-model organisms owing to several international initiatives such as the Earth BioGenome project (EBP), Darwin Tree of Life Project, The Vertebrate Genomes Project, 1000 Fungal Genomes Project and ERGA (European Reference Genome Atlas), among others. However, to answer ecological and evolutionary relevant questions we need not only the complete sequence of genomes but also accurate annotations including regulatory genes such as lncRNAs.

Most studies related to lncRNA identification rely on only one method of coding potential detection (Boltaña et al., 2016; Dettleff et al., 2020; Mu et al., 2016; Paneru et al., 2018; J. Quan et al., 2020; Ren et al., 2020; H. Xu et al., 2019). In contrast, our results based on an alignment-dependent (CPC) and two alignment-free methods (CPAT, FEELnc), indicate that 92.0%, 51.7% and 90.2% lncRNAs detected by CPC, CPAT and FEELnc respectively were called as lncRNAs by at least two tools. Different tools revealed to have varying performances (Schneider et al., 2017; Wucher et al., 2017) and the three tools used in our study perform well in predicting lncRNAs also in other species (Duan et al., 2021). Thus, lncRNAs predicted using our pipeline could be reliable resources for future studies to investigate lncRNAs functions. Our pipeline was integrated as an automatic tool, which can assemble transcripts based on the sequence alignment data, detect the lncRNAs from the reference genome through CPC, CPAT and FEELnc, and classify the lincRNAs detected by at least two methods. Hence, our pipeline is easily applied to other studies involving RNA sequencing datasets to investigate into lncRNAs detection and analysis.

We identified a total of 9,209 putative lncRNAs, of which 49% were expressed in at least 75% of our *A. polyacanthus* brain samples. While the numbers found in mammal species are higher (6,010 - 96,411 from the rhesus macaque to human; NONCODE v5.0, Fang et al., 2018) it has to be noted that in our case we only investigate one tissue, and the inclusion of more tissues is likely to increase the number of lncRNAs. Nonetheless, teleost model species show similar numbers of expressed lncRNAs. Zebrafish lncRNAs identified so far are 3,503 (Zhao et al., 2021) and between three- and six thousand lncRNAs have been found to be expressed across different tissues also in rainbow trout (Al-Tobasei et al., 2016), coho (Leiva et al., 2020) and Atlantic salmon (Boltaña et al., 2016). Other studies found higher numbers of lncRNAs in some teleost species, such as 31,984 in the brain of *Larimichthys crocea* (Liu et al., 2018) using CPC or 14,614 across several tissues of *Genypterus chilensis* (Dettleff et al., 2020) using CPAT. However, these studies only used one method to evaluate coding potential and did not ensure that all transcripts of a lncRNA were non-coding, which are less stringent criteria compared to what was applied here.

Overall, the majority of lncRNAs we identified throughout the genome of *A. polyacanthus* had relatively low expression. This observation is in accordance with many previous studies that report a typical low expression of lncRNAs across species and tissues (Guo et al., 2020; Hezroni et al., 2015; Necsulea et al., 2014; Quinn et al., 2016), as lncRNAs are constantly submitted to various regulation mechanisms and they are typically short-lived (Necsulea et al., 2014; Wu et al., 2014). Despite low expression levels, these lncRNAs can be functional since some regulatory mechanisms do not require high concentration of effector molecules (Aprea & Calegari, 2015). However, some lncRNAs can also be expressed at elevated levels and highly expressed intergenic lncRNAs (lincRNAs), for instance, are associated with diseases (Wan et al., 2013; F. Yang et al., 2011). In the brains of our coral reef fish, we also found some highly expressed lincRNAs with neighbouring coding genes involved in fundamental functions such as a GABAA receptor, circadian rhythm genes and genes related to signal transduction (such as DRD1L and ADA2B), important for normal brain function and neuronal activity (Bhat et al., 2010; Logan & McClung, 2019). In humans, rodents, and zebrafish, a lncRNA promotes the expression of homeobox transcription factors required for the development of GABAergic neurons Y-Aminobutyric acid (or GABA) which is the main inhibitory neurotransmitter in the vertebrate brain (Feng et al., 2006). Furthermore, lncRNAs have been shown to regulate core circadian rhythm genes in mammals (Mosig & Kojima, 2021). With a substantial fraction of lncRNAs affecting the gene expression of their neighbouring coding genes (Engreitz et al., 2016), it is no surprise to see elevated expression in lncRNAs neighbouring GABA_A_ receptor and core circadian rhythm genes as these are known to play important roles in the transcriptional response to ocean acidification (Kang et al., 2022; Schunter et al., 2021, 2016). As our RNAseq data is from ocean acidification experiments, this may suggest a regulatory involvement of lncRNAs in these key functions. Hence, our pipeline of lncRNA annotation and expression analysis allows for the discovery of new lncRNAs that may be involved in the response to an environmental change in a wild coral reef fish, but it also recovers conserved lncRNAs found to possibly be involved in essential functions also in other species.

### Functional responses of lncRNAs to environmental change

Our pipeline facilitates the investigation of lncRNAs as a potential regulatory mechanism in response to environmental change and opens doors to fields like environmental science and ecology. In our case study, we made use of a published RNA sequencing set of *Acanthochromis polyacanthus* brains from a volcanic CO_2_ seep, representing future ocean acidification conditions, and a control site. We detected differentially expressed lincRNAs and mRNAs in the brain of *A. polyacanthus* as a response to natural environmental differences (CO_2_ levels). Interestingly, for the coding genes that have a neighbouring lincRNA in close proximity, we found larger expression differences between fish from CO_2_ seeps and control sites suggesting a potential regulation of gene expression by neighbouring lncRNAs with the exposure to elevated CO_2_. For the differentially expressed lincRNAs, the set of neighbouring protein coding genes that were also differentially expressed are involved in functions related to pH regulation in fish. When fish respond to ocean acidification, carbonic anhydrase (CAHZ) is often upregulated, as in our data here, to catalyze the hydration of CO_2_ and play an essential role in acid-base and ion regulatory functions (Heuer & Grosell, 2014). A further reduction of intracellular pH in fish is necessary to prevent acidosis when faced with elevated CO_2_ levels in fish (Schmidt, 2019), and this process performed by coding gene GRIN2C may potentially be regulated by differentially expressed lincRNAs.

Further biological processes are known to be involved in the response to ocean acidification in fish brains. One of them is synaptic transmission and the downregulation of a lincRNA and the neighbouring coding gene CACNA1E (voltage-gated calcium channel complex) can contribute to synaptic transmission (Berecki et al., 2014), which was also found to behave similarly to a previous RNAseq *A. polyacanthus* data (Schunter et al., 2018). Some lincRNAs may trans-regulate the expression of ion transporters, such as calcium-activated potassium channel subunit alpha-1 (KCMA1), sodium leak channel non-selective (NALCN), and voltage-dependent P Q-type calcium channel subunit alpha-1A (CACNA1A), which play critical roles in the neural signal transduction (Gadsby, 2009) and were also reported in our previous study (Kang et al., 2022). In addition, immune responses are commonly associated with ocean acidification in fish including our study species (De Souza et al., 2014; Kang et al., 2022; Machado et al., 2020), and here we find differential expression of lincRNAs neighbouring and possibly trans-regulating a variety of immune response genes that show co-expression in a gene network significantly correlated with CO_2_ levels. This suggests that lincRNAs annotated in our coral reef fish show differential expression in response to an environmental factor, CO_2_ levels, and that some of these differentially expressed lincRNAs are in close proximity to coding genes that are known to have functional implications in the response to ocean acidification in fishes.

We identified candidate lincRNAs to be involved in acclimation to elevated CO_2_ levels and future genetic manipulation experiments would allow us to verify the direct effect of a lincRNA on the nearby coding gene (Akay et al., 2019; Rodriguez-Lopez et al., 2022). CRISPR/Cas9 based genome editing is a promising technique to experimentally evaluate the role of lincRNAs, although it is not readily available for most non-model organisms. Our study is a start to understand regulatory elements potentially involved in the regulation of the gene expression patterns observed with an environmental change. This case study presents a pipeline that can be applied by using RNA sequencing reads, now frequently incorporated into experimental studies, and adds an epigenetic mechanistic aspect with no additional sequencing effort needed. We demonstrate that lncRNAs can be easily annotated de novo in non-model organisms and encourage more molecular ecologists to discover lncRNAs in their study species and investigate the mechanistic role of lncRNA in response to environmental change.

## ACKNOWLEDGMENTS

We would like to thank Timothy Ravasi and Philip L. Munday for for helping with the collection and sequencing of the large RNA sequencing datasets produced in previous studies. We are grateful to Helen Leung, Ho Wu Cheuck and Kam Yan Chit for their support with this project.

## DATA AVAILABILITY

The RNA sequencing raw data used in this study is found in the Bioprojects: PRJNA691990; PRJNA311159; PRJNA658203 (Reviewer link as the last is not published yet: https://dataview.ncbi.nlm.nih.gov/object/PRJNA658203?reviewer=47r5c4kubkjn124c770t5vvt60)

The lncRNA annotation of *Acanthochromis polyacanthus* is available here: 10.6084/m9.figshare.20045780. Reviewer link: https://figshare.com/s/33802daaa28dc7ccd876

## CODE AVALABILITY

The scripts have been deposited in Github (https://github.com/jinglkang/lncRNAs_detect).

## SUPPLEMENTARY FIGURES

**Figure S1:**
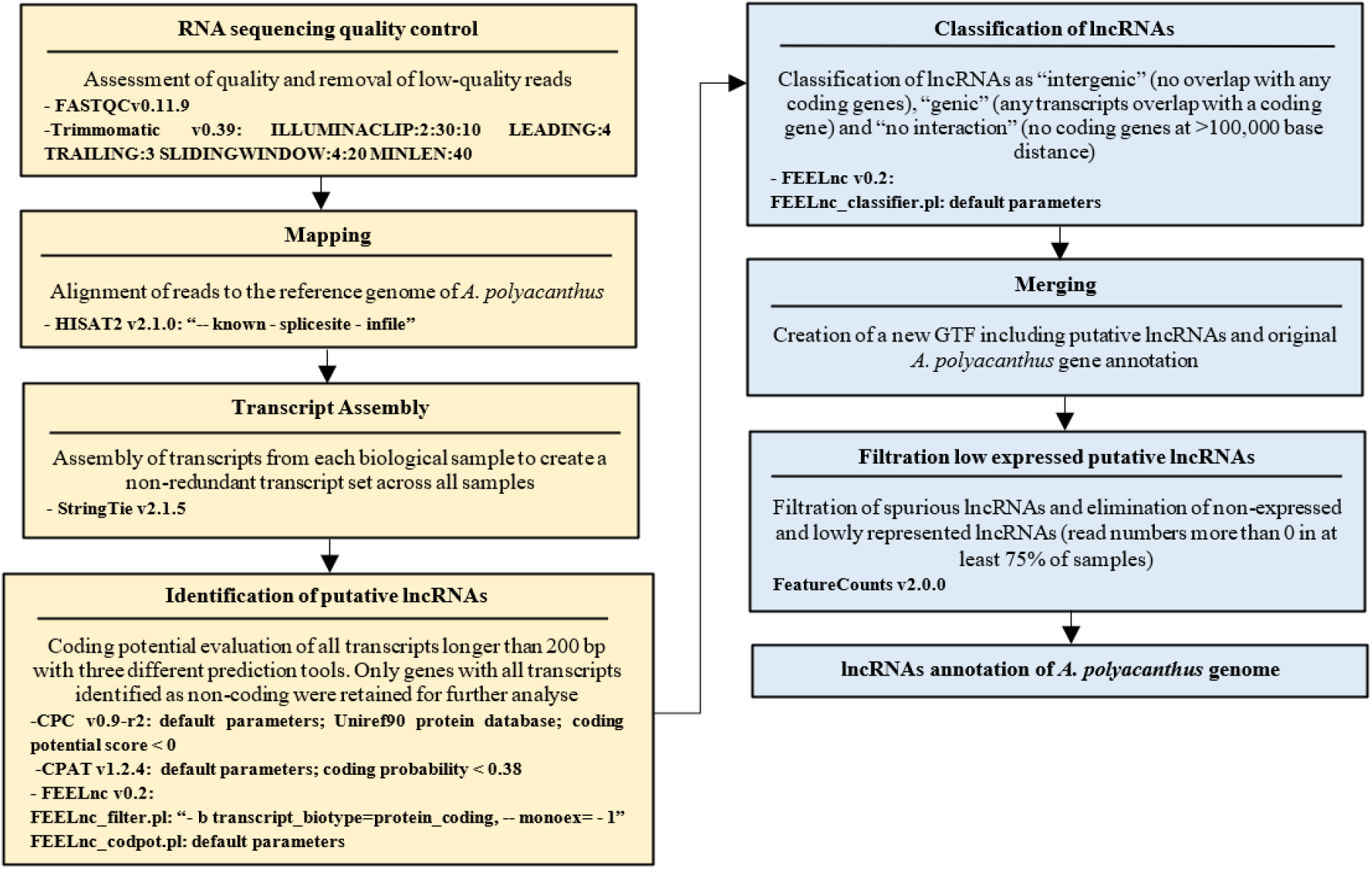
Detailed flowchart illustrating the process for long non-coding RNA (lncRNA) identification including the programs and parameters used

**Figure S2:**
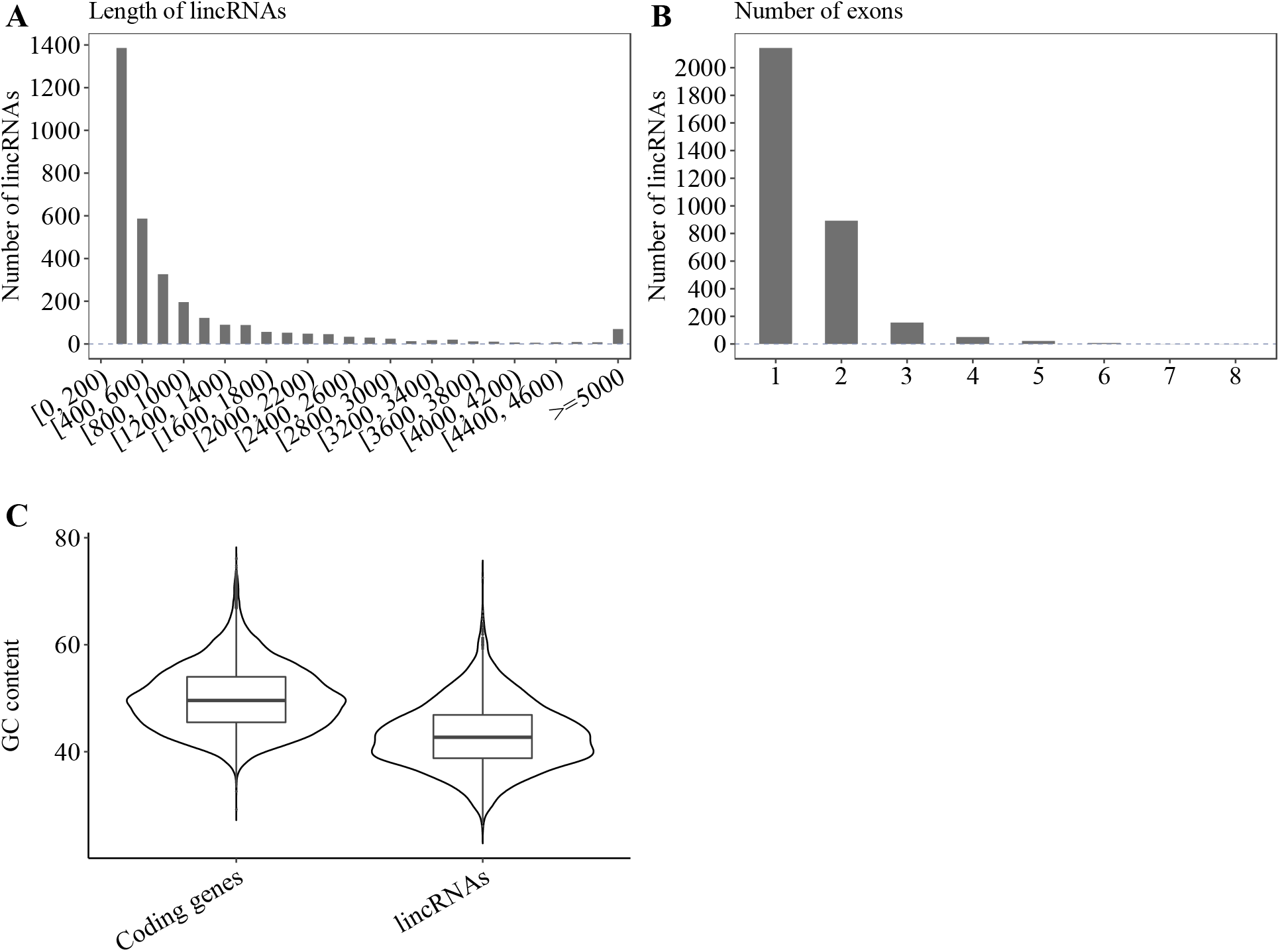
Sequence features of intergenic long non-coding RNAs (lincRNAs) in *Acanthochromis polyacanthus*. A. Length distributions of lincRNAs. B. Distributions of the number of exons in lincRNAs. C. The GC content in coding genes and lincRNAs. The GC content of lincRNAs were significantly lower (two-sample wilcoxon rank sum test, p < 2.2e-16) than coding genes.

**Figure S3:**
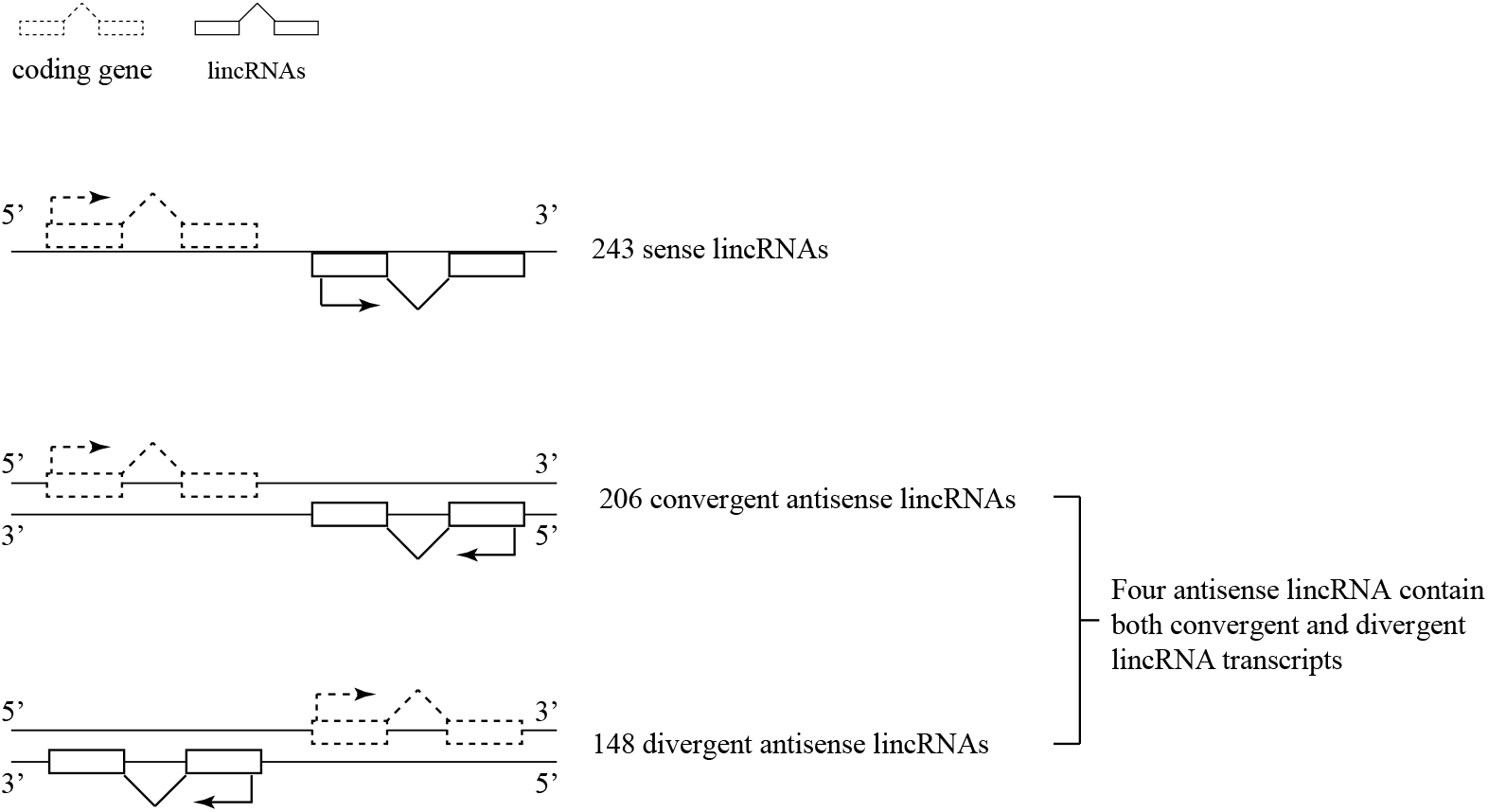
The orientation of intergenic long non-coding RNAs (lincRNAs) in Acanthochromis polyacanthus.

**Figure S4:**
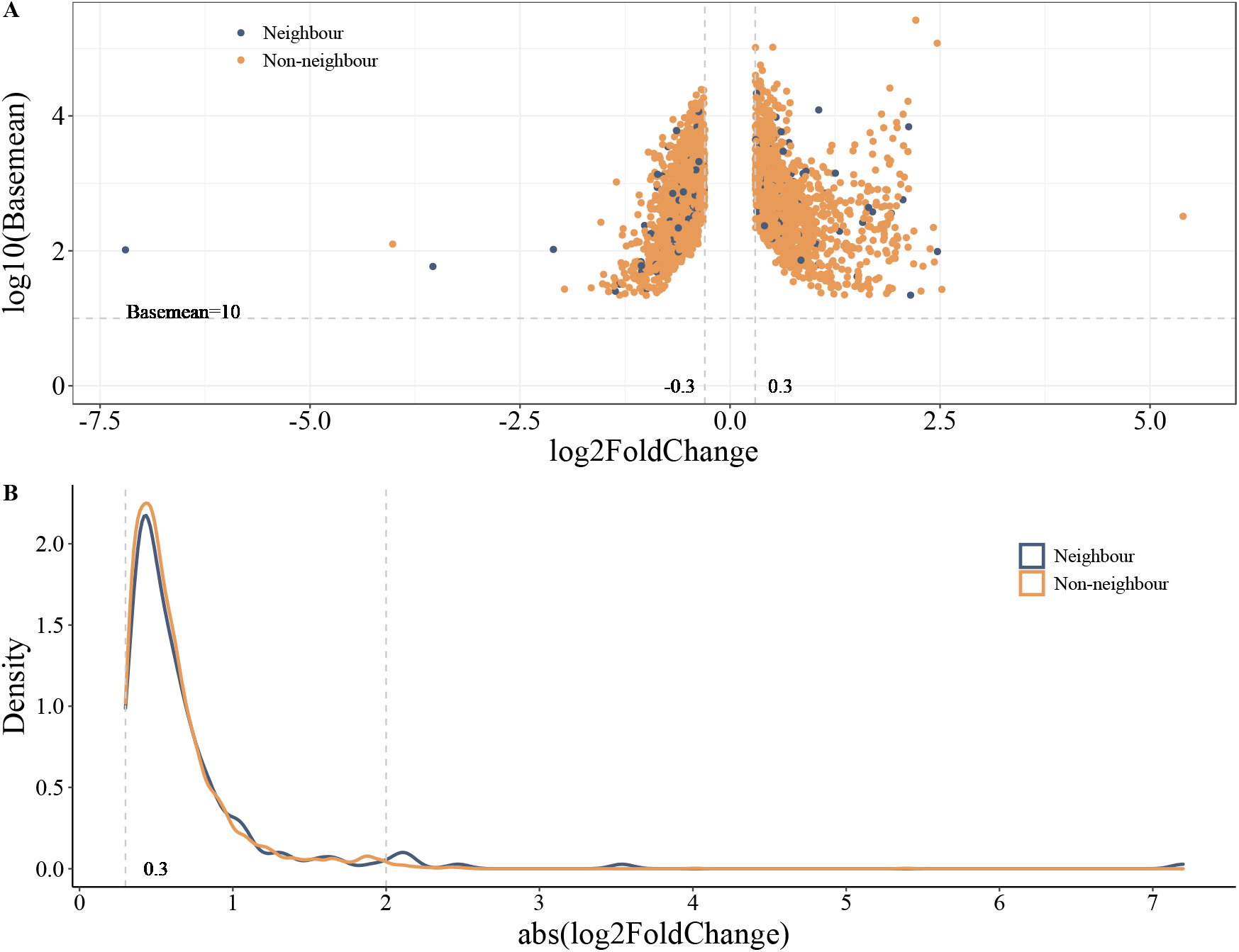
Expression pattern comparison between coding genes with (blue) and without (orange) neighbouring lincRNAs. A. Volcano plot of all expressed coding genes. The gene expression log2 fold change and log10(basemean) between samples from CO_2_ seep and control site are reported on the X and the Y axes, respectively. B. Density of log2FoldChange absolute value of gene expression between samples from CO_2_ seep and control site.

